# scOLAR: Ontology-Anchored Open-Set Annotation of Single-Cell RNA-seq Data

**DOI:** 10.64898/2026.09.05.749179

**Authors:** Yuqiao Liu, Siyu Yi, Hengchuang Yin, Wei Ju

## Abstract

Single-cell RNA sequencing profiles cellular heterogeneity at atlas scale, making automated annotation essential. However, target datasets often contain novel cell types missing from incomplete references. We present scOLAR, an ontology-guided open-set framework that learns prototypes over the Cell Ontology and uses both reference and target expression to annotate known classes while detecting unfamiliar populations. Guided by ontology hierarchies and decision-boundary regularization, scOLAR penalizes coarse-lineage misclassification and groups novel cells without requiring predefined cluster counts. Across benchmarks, scOLAR achieves a novelty-detection AUROC of 0.9726 and an average precision of 0.9871, enabling structured post-hoc lineage-level interpretation of populations absent from the reference.

## 1 Background

Single-cell RNA sequencing (scRNA-seq) provides a detailed view of the cellular composition of tissues and how it changes during development and disease [1, 2]. Large initiatives such as the Human Cell Atlas [3], Tabula Muris [4] and developmental atlases [5] provide reference data that connect gene expression profiles to cell types. These resources allow new experiments to be compared with previously characterised cell populations.

Reliable cell-type annotation is essential for these comparisons. Seurat [6] and Scanpy [7] support large-scale analysis and clustering, but assigning biological identities still requires marker information [8] and expert judgement [9, 10]. As datasets grow, manual review of every cluster becomes increasingly demanding. Computational methods reduce this burden by transferring cell-type information from annotated reference data to new datasets [11]. Neural classifiers, generative models and methods that combine annotation with clustering provide different ways to make use of this information [12–17].

In practice, reference-based annotation depends on how well the reference represents the new dataset. Differences in tissue, developmental stage and disease context affect reference coverage, and rare populations may be missed during sampling [3, 5, 9]. A new dataset, referred to here as the target, can therefore contain cell types with no corresponding reference label. Assigning these cells to an existing type can obscure populations that deserve further investigation. This creates two related tasks, recognising known cell types and identifying cells that may represent populations absent from the reference. Throughout this study, a novel type is one that is absent from the reference.

Open-set annotation allows a model to leave cells unassigned when the available labels do not adequately describe them [18]. MARS learns cell-type landmarks to identify unfamiliar populations [19]. scNym, scArches and ItClust use different forms of transfer learning to adapt reference information to target data [20–22]. Other approaches use spatial context [23], multiple references [24] or pathway knowledge [25] to support annotation. OVAAnno addresses novel-type detection in single-cell chromatin accessibility data and also includes evaluation on scRNA-seq data [26]. When a specific assignment is uncertain, hierarchical rejection can retain a broader cell-type label [27].

Unassigned cells also need to be organised into groups that can be examined biologically, and the number of these groups is usually unknown. This connects open-set annotation with generalised category discovery, in which labelled examples support recognition of known classes and discovery of new ones [28, 29]. Recent single-cell frameworks combine these tasks, including scGAD, scEVOLVE and scBOL [30–34]. In incomplete-reference settings, this means annotating known cells while organising unfamiliar cells without target labels or prior knowledge of how many novel types are present. These groups also need biological context so that researchers can decide which populations to examine further.

The Cell Ontology (CL) provides a shared vocabulary for cell types and records relationships between them [35, 36]. It links specific cell types to broader categories and includes developmental relationships. This can be useful when the data support a broad lineage but not a specific cell type. OnClass uses the ontology to predict cell types with no labelled training examples [37]. Other approaches use it to keep predictions consistent across levels of the hierarchy [38] or to guide language models towards standard cell-type names [39].

Here we introduce scOLAR to annotate known cell types, identify unfamiliar cells and organise them for biological follow-up. The model learns a shared representation from labelled reference cells and unlabelled target expression profiles. Relationships in the CL guide how reference cell types are represented, helping distinguish confidence in a specific type from similarity to a broader lineage. A threshold calibrated using reference cells alone flags target cells as potentially novel. The model assigns reference labels to the remaining cells and clusters the flagged cells without requiring the number of novel types in advance. For novel groups that can be linked to the ontology with sufficient confidence, a separate step suggests broader cell categories. These categories can then be considered together with marker expression during biological review.

Across five scRNA-seq benchmarks, scOLAR separates known and novel cells with an area under the receiver operating characteristic curve of 0.9726 and an average precision of 0.9871. The resulting novel-cell groups can then be examined using marker expression together with their ontology-based descriptions.

## 2 Results

### 2.1 Integrated workflow and evaluation setting

Annotation with an incomplete reference requires several connected decisions. The model needs to recognise known cell types, flag unfamiliar cells and group those cells for biological review. scOLAR combines these steps in one workflow (Figure 1). Training uses labelled reference cells and unlabelled target expression profiles. The prototype head contains one learnable row for every term in the compiled CL. The rows mapped to source classes define the labels that can be assigned to known cells.

**Fig. 1.**
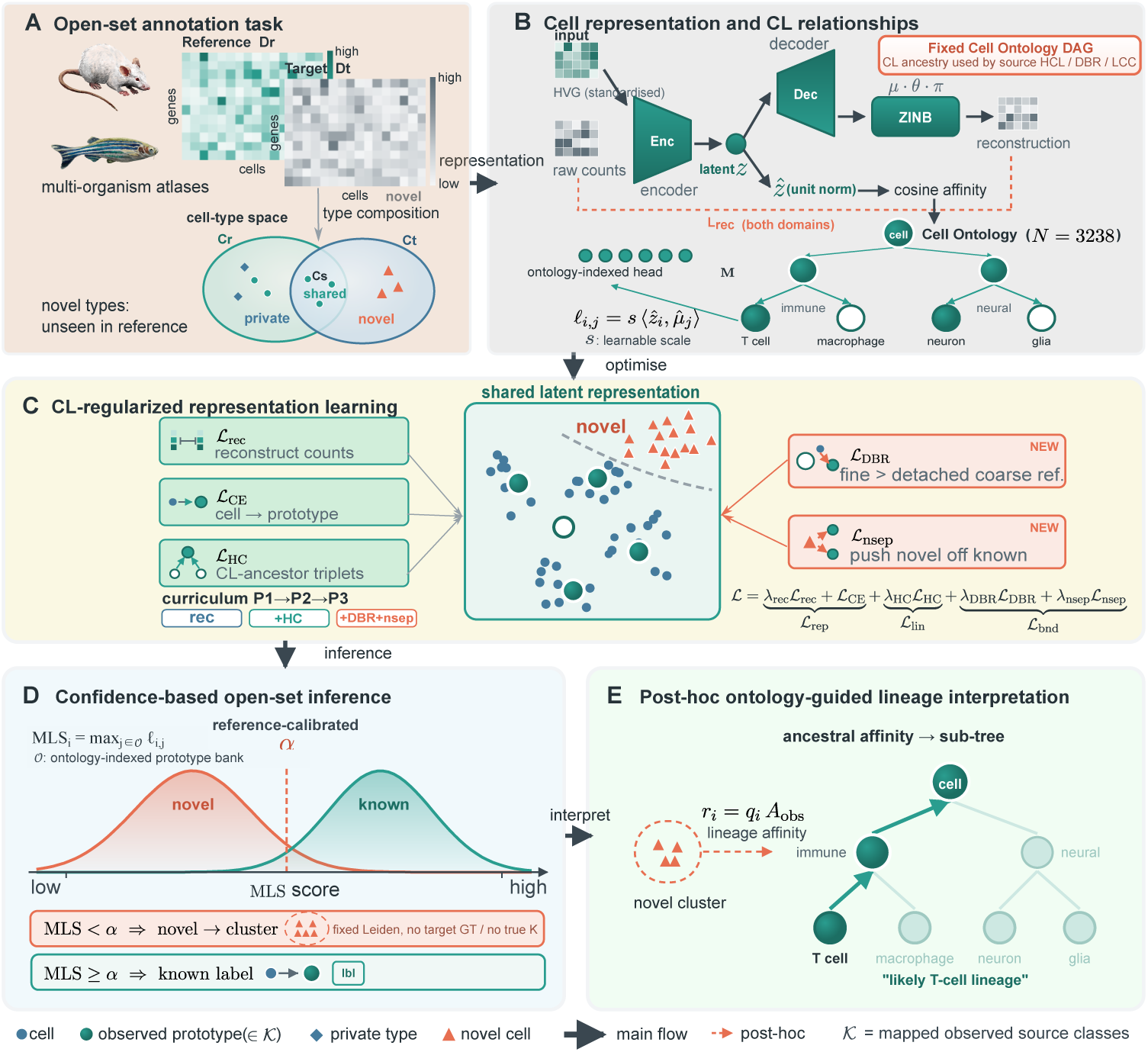
Architecture and inference workflow of scOLAR. **(A)** Training uses a labelled reference and unlabelled target expression. **(B)** A shared encoder learns cell representations, and each row of the prototype head corresponds to a Cell Ontology term. The diagram highlights the mapped source-class rows used to assign known labels. **(C)** Objectives based on Cell Ontology relationships guide source prototype geometry and the separation of fine-type and broad-lineage evidence. Reconstruction and novel separation also contribute to training. **(D)** The maximum logit over the complete head is compared with a reference-calibrated threshold to assign each cell to Known or PredNovel. Known cells receive the source-class label with the highest score. Leiden groups all PredNovel cells without a supplied *K*. **(E)** The Lineage Consistency Check uses source-class affinities and Cell Ontology ancestry to describe eligible communities after clustering.

For each target cell, scOLAR compares the maximum logit over the complete head with a threshold calibrated from reference cells. The two resulting branches are called Known and PredNovel. Cells in Known receive a source-class label. Leiden then partitions all PredNovel cells without a predefined number of novel populations. A separate analysis uses the ontology to describe the lineage affinities of eligible communities (Figure 1). We assess each step separately. Ranking measures the continuous novelty score, routing is the threshold-based Known/PredNovel decision, classification assigns source labels within Known, and discovery groups cells within PredNovel.

We evaluate the workflow on five established open-set scRNA-seq benchmarks, Cao [40], Quake 10x and Quake Smart-seq2 [4], Wagner [41], and Zeisel 2018 [42]. Shared, reference-private and target-private cell types follow the published benchmark protocol [34]. Together with the earlier cross-dataset analyses, the dataset collection covers four organisms, multiple tissues and five sequencing platforms (Table S1 in Additional file 1). Each primary summary averages four independent runs. Training uses unlabelled target expression, a setting known as transductive learning. Target labels and the number of target-private classes are withheld. The epoch-100 checkpoint, reference-calibrated threshold and fixed Leiden resolution are set before target scoring. Ground-truth labels enter only after outputs are fixed. This dataset range supports evaluation across several settings, although general cross-species validity remains to be established.

### 2.2 Benchmark context and common-evaluator comparison

Published benchmarks provide context for the tasks and datasets considered here. Table 1 includes seven earlier methods, among them scCNC [43] and scDECL [44]. Their results use different training information, selection rules, replication and deployment assumptions. We retain these historical values to show the earlier evidence. They cannot support a newly controlled ranking of all eight methods.

**Table 1.** Published benchmark results for the Cao and Quake datasets.

| Method | Cao |  |  | Quake_10x |  |  | Quake_Smart-seq2 |  |  |
| --- | --- | --- | --- | --- | --- | --- | --- | --- | --- |
|  | Known | Novel | Overall | Known | Novel | Overall | Known | Novel | Overall |
| scCNC | 55.8 | 42.8 | 28.7 | 84.4 | 65.3 | 68.5 | 58.0 | 35.3 | 38.5 |
| scDECL | 51.4 | 41.6 | 26.3 | 31.7 | 45.9 | 26.1 | 22.0 | 30.3 | 25.9 |
| MARS | 92.7 | 58.2 | 63.1 | 96.4 | 49.8 | 67.5 | 88.1 | 78.9 | 78.3 |
| ItClust | 3.4 | 45.0 | 48.0 | 54.3 | 43.3 | 53.2 | 10.9 | 62.0 | 65.9 |
| scNym | 98.5 | 63.1 | 61.2 | 98.5 | 48.1 | 53.7 | 95.3 | 69.6 | 65.9 |
| scArches | 78.0 | 45.4 | 57.7 | 90.3 | 57.3 | 70.1 | 64.0 | 56.1 | 58.1 |
| scBOL | 96.5 | 74.6 | 77.4 | 98.0 | 65.8 | 77.5 | 96.2 | 82.1 | 82.5 |
| **scOLAR (ours)** | $99.2 \pm 0.2$ | $78.1 \pm 1.7$ | $89.5 \pm 0.9$ | $99.3 \pm 0.3$ | $68.6 \pm 2.0$ | $87.2 \pm 1.0$ | $97.8 \pm 0.1$ | $78.3 \pm 1.2$ | $88.3 \pm 0.7$ |

**Table 1 Published benchmark context (continued).**
| Method | Wagner |  |  | Zeisel.2018 |  |  |
| --- | --- | --- | --- | --- | --- | --- |
|  | Known | Novel | Overall | Known | Novel | Overall |
| scCNC | 83.1 | 57.4 | 59.8 | 64.9 | 79.1 | 70.1 |
| scDECL | 32.5 | 48.8 | 35.0 | 55.8 | 69.4 | 49.5 |
| MARS | 78.0 | 53.0 | 54.8 | 89.8 | 87.0 | 83.9 |
| ItClust | 29.4 | 30.8 | 36.0 | 32.1 | 69.3 | 63.7 |
| scNym | 93.9 | 44.6 | 43.5 | 99.3 | 62.2 | 62.4 |
| scArches | 65.2 | 41.0 | 46.8 | 73.1 | 63.2 | 63.6 |
| scBOL | 94.9 | 54.7 | 62.5 | 96.5 | 91.7 | 89.1 |
| **scOLAR (ours)** | **99.9 ± 0.0** | **58.0 ± 2.5** | **72.1 ± 1.6** | **99.6 ± 0.1** | **99.3 ± 0.1** | **99.4 ± 0.1** |

To compare outputs under shared scoring rules, we evaluate scOLAR and strict scBOL with a common downstream evaluator (Figure 2). Known measures classification accuracy on ground-truth-known target cells. Novel weighted, Novel macro and adjusted Rand index (ARI) measure recovery on ground-truth-novel cells. For this benchmark analysis, clustering receives the true number of novel classes only after training. OverallJ gives the joint benchmark result. The methods retain their own training assumptions. Strict scBOL receives the true target class count during training, an oracle input that scOLAR does not use. For scOLAR, true *K* enters only the post-training benchmark clustering step.

**Fig. 2.**
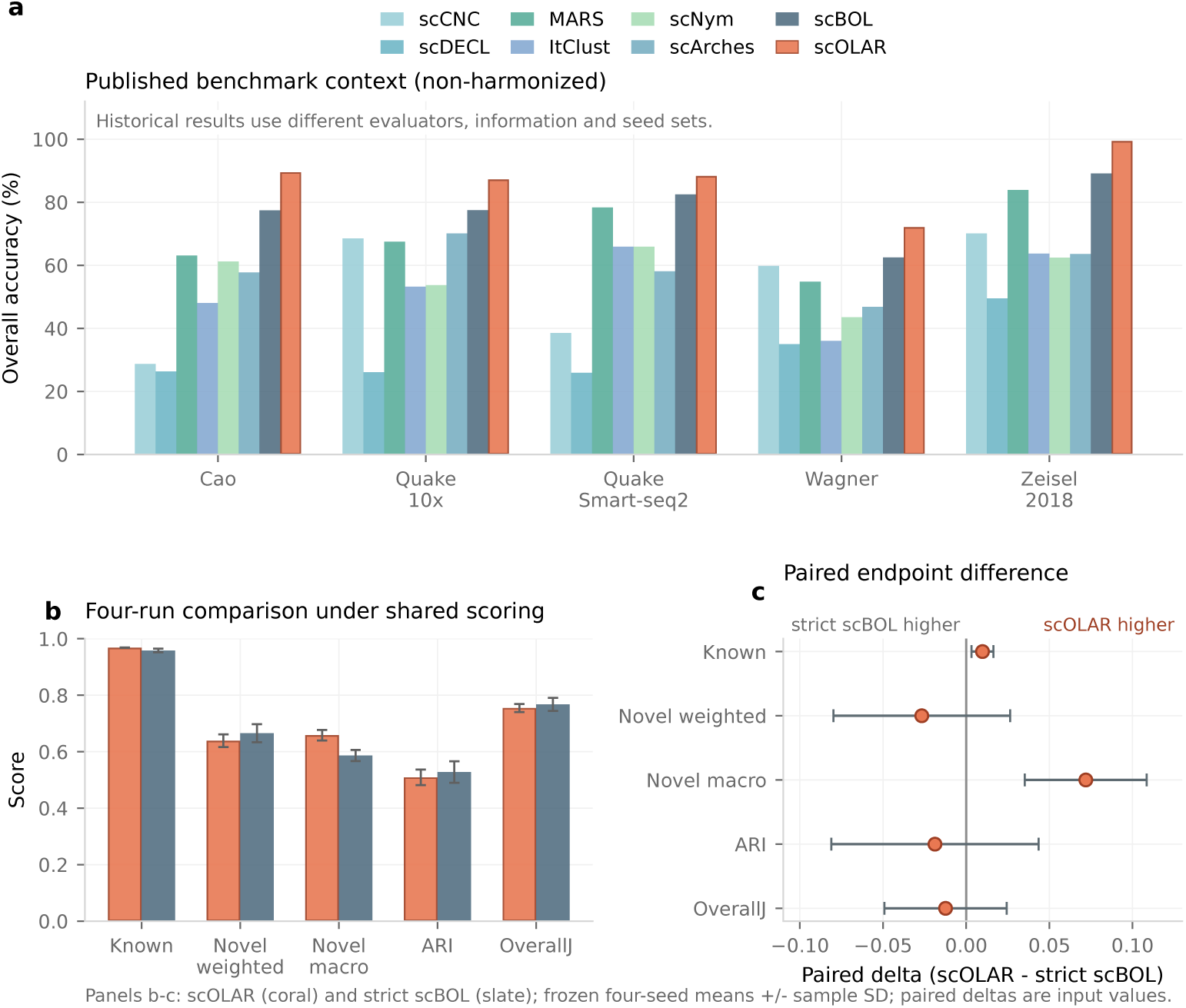
Benchmark context and common-evaluator performance of scOLAR. **(a)** Published Overall values summarise the historical results in Table 1, whose evaluation settings differ across methods. **(b)** Means under the common evaluator compare scOLAR with strict scBOL for Known, Novel weighted, Novel macro, ARI and OverallJ. **(c)** Paired differences are scOLAR minus strict scBOL. Positive values favour scOLAR for the metric shown. Error bars show sample standard deviations across four independent runs (*n* = 4). Ground-truth-novel membership and true *K* enter benchmark clustering and scoring only after predictions are fixed. Strict scBOL receives the true target class count during training. The common evaluator therefore compares outputs under shared scoring rules, with the training information and deployment assumptions of each method preserved.

The methods perform differently across metrics. scOLAR has higher Known accuracy (0.9682 ± 0.0006 versus 0.9584 ± 0.0064, paired difference +0.0098 ± 0.0065) and higher Novel macro recovery (0.6586 ± 0.0187 versus 0.5867 ± 0.0197, difference +0.0720 ± 0.0367). Strict scBOL has higher Novel weighted accuracy (0.6656 ± 0.0319 versus 0.6389 ± 0.0225), ARI (0.5280 ± 0.0382 versus 0.5093 ± 0.0277) and OverallJ (0.7673 ± 0.0232 versus 0.7549 ± 0.0143). Section S5 and Table S5 in Additional file 1 report all values and paired differences. scOLAR therefore recovers novel types more accurately when each type receives equal weight. Measures weighted by cell abundance and global clustering measures favour strict scBOL. The results support these specific strengths and trade-offs, with no overall advantage across all metrics.

### 2.3 Novelty detection using reference-based calibration

A practical annotation method needs to recognise unfamiliar cells before their labels or number are known. We assess this ability using scOLAR’s own PredNovel decisions (Figure 3). This deployment analysis starts with all cells the model flags, without first selecting ground-truth-novel cells or supplying their true count. Across five datasets and four independent runs, the fraction of target-private cells is 0.6564 ± 0.0018. The maximum logit score (MLS) distinguishes known and novel cells with an area under the receiver operating characteristic curve (AUROC) of 0.9726 ± 0.0012 and average precision of 0.9871 ± 0.0006. At the threshold calibrated from reference cells, novelty precision is 0.8837 ± 0.0052, recall is 0.9782 ± 0.0015, and F1 is 0.9284 ± 0.0023. The PredNovel fraction is 0.7273±0.0062, which exceeds the actual target-private fraction. This operating point favours sending potentially unfamiliar cells for further analysis.

**Fig. 3.**
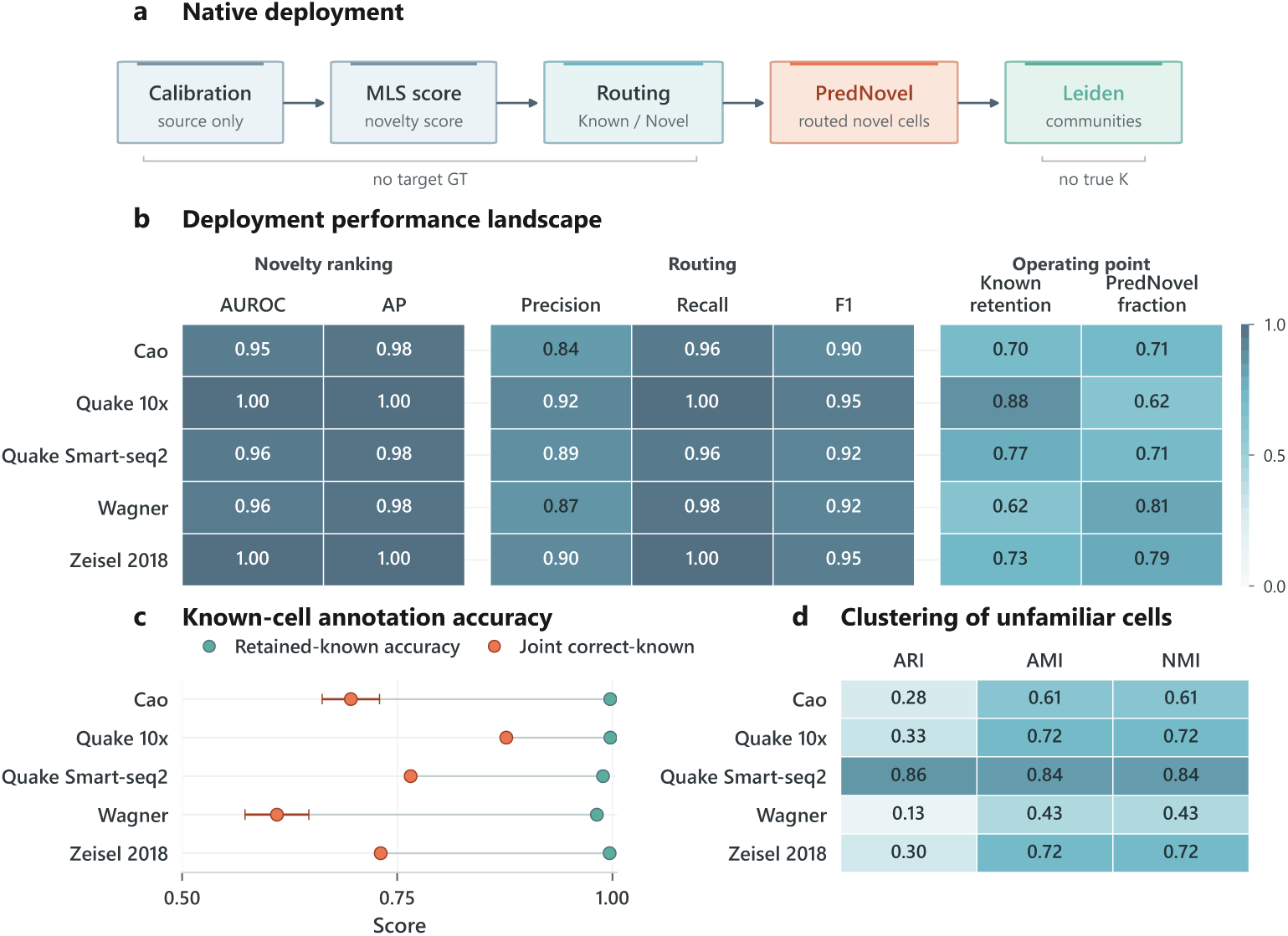
Deployment performance and discovery without a predefined number of novel types. **(a)** The workflow uses maximum-logit ranking and a threshold calibrated from reference cells to classify known cells and group unfamiliar cells with Leiden, without a supplied *K*. **(b)** Heat maps show novelty ranking and threshold-based decisions for each dataset. **(c)** Paired run-level points show conditional accuracy among retained known cells and the joint correct-known rate. Known retention is not plotted. The horizontal axis spans 0.5–1.0. **(d)** The heat map reports clustering quality on the fixed partition of all PredNovel cells. Table 2 gives the aggregate values. PredNovel includes every cell sent to the novel branch, including errors. Deployment decisions use no target labels or true *K*.

**Table 2.**
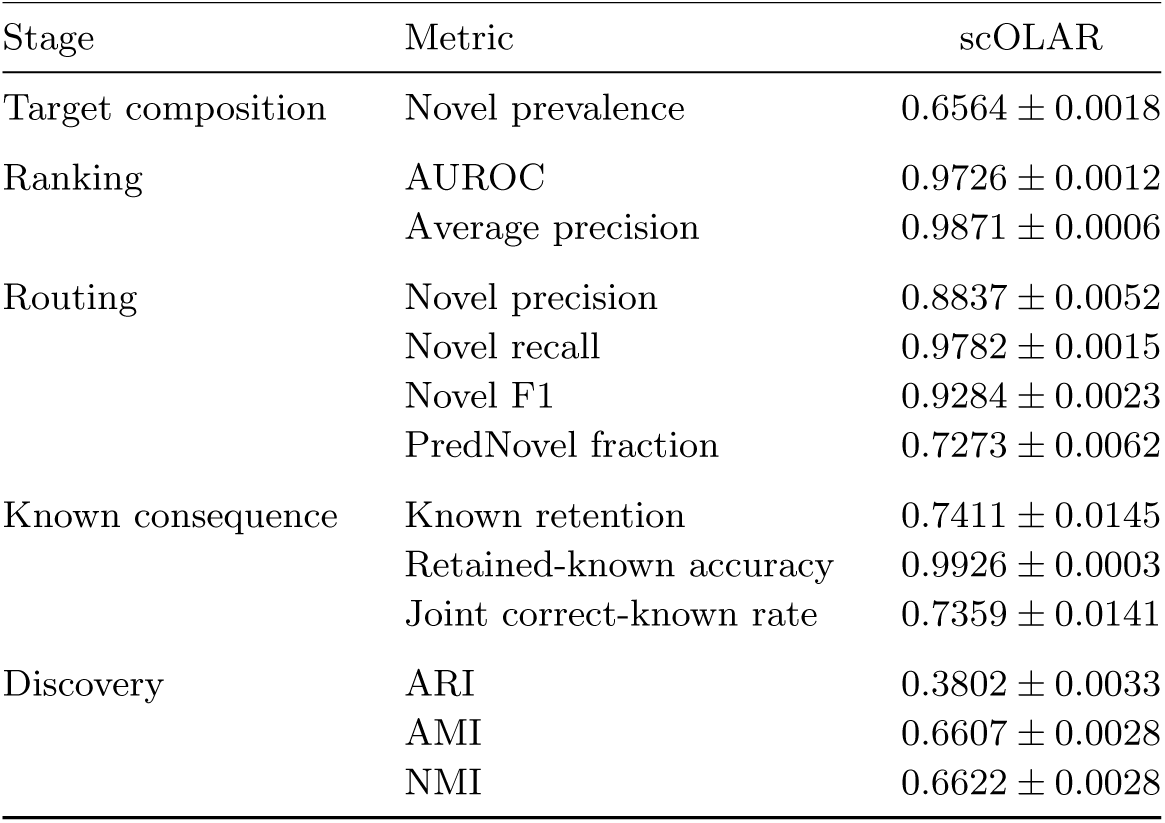
scOLAR deployment summary. Values are equal-dataset means *±* sample standard deviations across four independent runs (*n* = 4). PredNovel includes all cells the model sends to the novel branch. Known retention is the fraction of ground-truth-known cells retained as Known. Retained-known accuracy is conditional on this subset. Joint correct-known measures the fraction of known cells that are both retained and correctly labelled, calculated directly for individual cells. Discovery evaluates the fixed Leiden partition of all PredNovel cells, formed without target labels or true *K*. Prevalence is the actual target-private fraction and provides context for the threshold-based decisions.

This conservative decision also sends some known cells to the novel branch. Known retention is 0.7411 ± 0.0145. Among the retained known cells, classification accuracy is 0.9926 ± 0.0003, and the joint rate of retaining a known cell and assigning its correct label is 0.7359±0.0141. Most of the reduction in known-cell performance thus occurs at the known/novel decision. Classification errors among retained cells contribute much less. Table 2 summarises deployment performance. Tables S6 and S7 in Additional file 1 give the deployment and end-to-end known-cell summaries across datasets. The joint correct-known rate is calculated directly for individual cells. Multiplying the aggregate retention and conditional accuracy would give a different quantity.

### 2.4 Discovery without a predefined number of novel types

Cells flagged as unfamiliar need to be grouped without advance knowledge of how many novel types are present. We apply Leiden once at fixed resolution 1.0 to all PredNovel cells. Neither target labels, a ground-truth-novel filter nor the number of target-private classes enters clustering. The resulting partition has ARI 0.3802 ± 0.0033, adjusted mutual information (AMI) 0.6607 ± 0.0028 and normalised mutual information (NMI) 0.6622 ± 0.0028 (Figure 3d and Table 2). These scores evaluate the population that scOLAR sends to discovery, including cells assigned to this branch in error. They therefore measure the combined effect of the known/novel decision and subsequent clustering.

### 2.5 Exploratory metric disagreement in Quake Smart-seq2

Averaging recovery over cells or over cell types gives different weight to abundant populations. To understand the disagreement on Quake Smart-seq2, we examined class abundance and local geometry after seeing the aggregate results (Figure 4). scOLAR’s Hungarian-matched Novel weighted accuracy is 0.6389 ± 0.0511, compared with 0.7172 ± 0.0374 for strict scBOL. Novel macro accuracy is higher for scOLAR at 0.6854 ± 0.0649, compared with 0.5272 ± 0.0395. Cluster purity, 15-nearest-neighbour agreement and same-label graph-edge agreement also remain high for scOLAR. Much of the weighted difference comes from numeric class 2, the most abundant novel class. It contains 4,394 cells, represents 30.33% of ground-truth-novel cells and contributes approximately −0.1114 to the four-run weighted-accuracy difference.

**Fig. 4.**
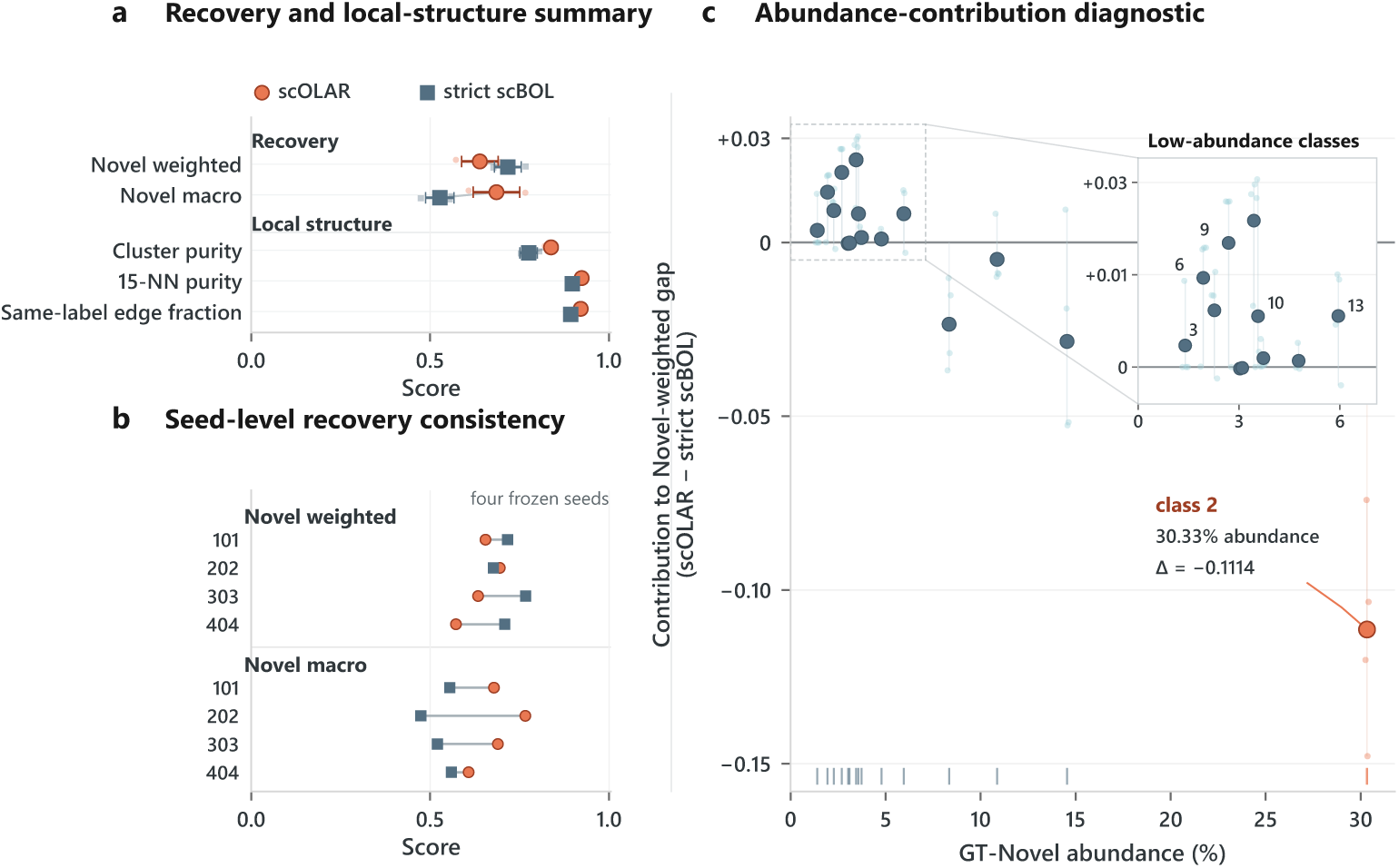
Exploratory analysis of metric disagreement in Quake Smart-seq2. **(a)** scOLAR has lower Novel weighted and higher Novel macro recovery, together with high local purity and neighbourhood agreement. Small pale points represent individual runs and large solid points their aggregate. **(b)** Run-level values show the direction of each recovery metric. **(c)** Numeric class 2 accounts for much of the weighted difference (*n* = 4,394, 30.33% of ground-truth-novel cells, mean contribution approximately *−*0.1114). The inset enlarges the low-abundance region of the same data and contains no fitted trend line. Numeric class identifiers are benchmark labels without an assigned biological identity in this analysis. The analysis follows the primary results and is descriptive. It does not establish a causal explanation.

A large class can dominate the cell-weighted result even when the average across classes favours the other method. This accounts for the direction of the two summaries. The analysis began after the primary metric pattern was known, so it serves to generate hypotheses. It identifies neither a biological mechanism nor grounds for changing the primary interpretation.

### 2.6 Ontology-guided interpretation of discovered communities

A group of unfamiliar cells needs biological context before it can be investigated further. The Lineage Consistency Check (LCC) provides this context after the PredNovel partition is fixed. It combines source-class affinities through CL ancestry and assigns the category Consistent to an eligible community when the predefined Top-3 coverage threshold is met. Communities that do not meet the threshold remain Ambiguous. Figure 5 illustrates the analysis on Zeisel 2018 using UMAP [45]. Marker expression supports an astrocyte-like description for one community, including expression of the canonical water-channel marker Aqp4 [46, 47].

**Fig. 5.**
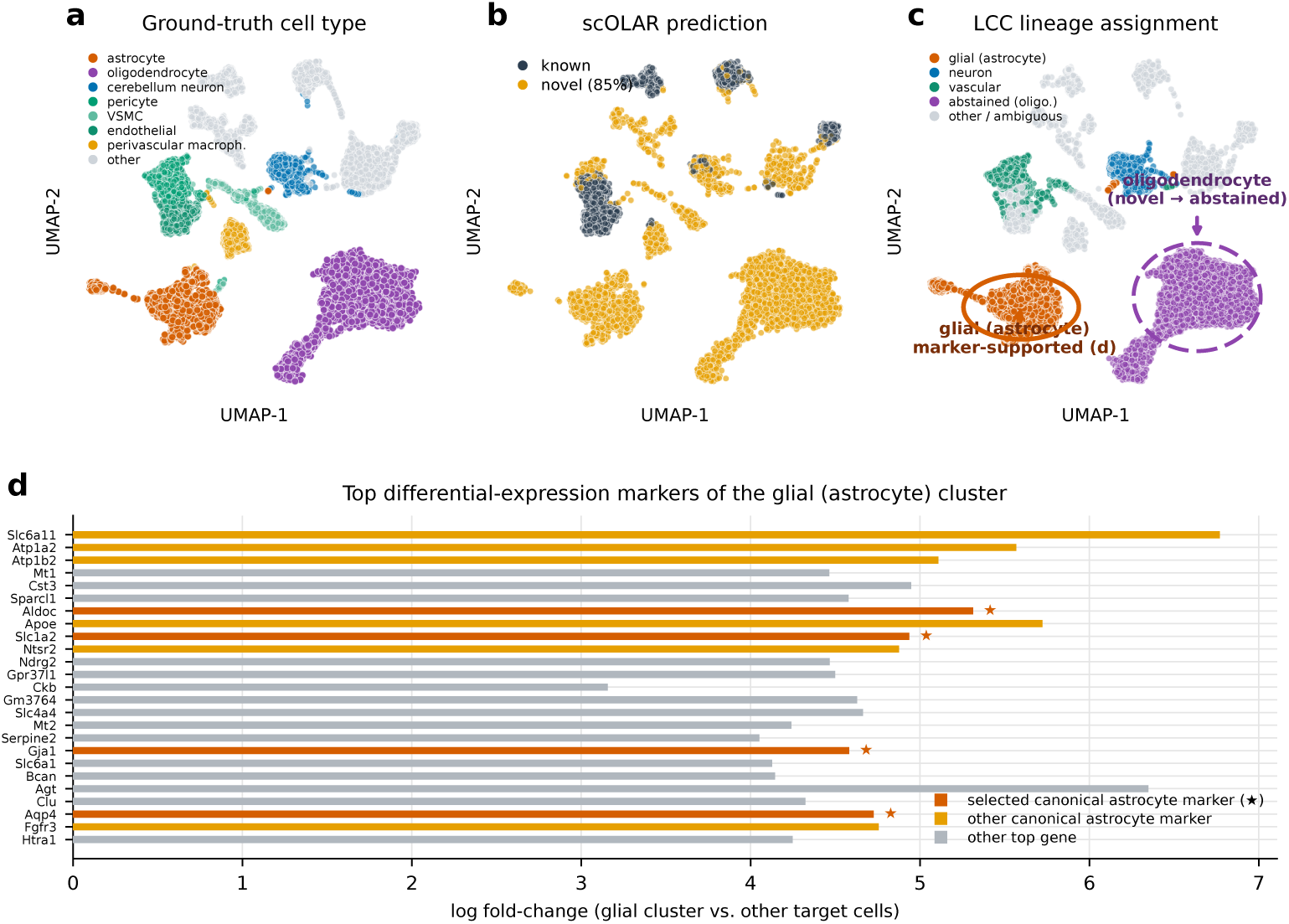
Example of ontology-guided interpretation of novel communities. (a–b) A Zeisel 2018 embedding shows the known/novel decisions and an already formed community. **(c)** LCC combines source-class affinities with Cell Ontology ancestry to describe that community. **(d)** Marker expression, including Aqp4, provides supporting biological context. This example comes from an earlier development run. Marker agreement alone does not independently validate LCC, the final model’s lineage descriptions or a causal benefit of ontology structure.

This example illustrates how a discovered community can be linked to a lineage hypothesis for review. It does not measure prospective LCC accuracy. The analysis covers only eligible communities, and its yield varies by dataset. The resulting profiles organise evidence for expert review. They leave the primary known/novel decisions and clustering unchanged and do not establish a new biological identity.

## 3 Discussion

When a reference is incomplete, cell-type annotation needs to accommodate unfamiliar populations and give those populations biological context. scOLAR connects these tasks in one workflow. CL relationships guide selected training objectives for reference cell types. A threshold calibrated from reference cells identifies potentially novel cells, Leiden groups them without a predefined number of populations, and LCC provides lineage descriptions for review. Each component has a specific role. MLS calibration, novel separation and Leiden use general open-set procedures. The ontology contributes to model indexing, the source HCL and DBR objectives, and the final descriptive analysis. Our conclusions concern this complete workflow. The prototype head does not constitute a semantic embedding of the entire Cell Ontology.

The comparison with strict scBOL shows that the methods have different strengths. scOLAR has higher accuracy on known cell types and higher average recovery across novel types. Giving each type equal weight is useful when small populations might otherwise contribute little to an overall score. Strict scBOL performs better on recovery weighted by cell abundance, ARI and OverallJ. The choice of metric therefore affects how the comparison is interpreted. The results also need to be read with the training conditions in mind, because strict scBOL receives the true target class count during training.

Deployment performance clarifies the practical cost of scOLAR’s novelty threshold. Novelty ranking is strong, and the reference-calibrated threshold gives high precision and recall. Approximately three quarters of ground-truth-known cells remain in the known branch. Those retained cells are labelled with high accuracy. The main cost is therefore the additional known cells sent for discovery and review. This conservative choice can reduce the risk of assigning unfamiliar populations to an existing type, but it increases the number of cells requiring follow-up. In practice, the preferred threshold depends on how an application weighs known-cell retention against the risk of overlooking unfamiliar populations.

The Quake Smart-seq2 analysis helps explain why class-balanced and cell-weighted recovery can point in different directions. One abundant numeric class accounts for much of the weighted deficit, even though the macro score and local neighbourhood summaries favour scOLAR. This pattern is consistent with the different weights assigned to cells and classes. It does not identify the class biologically or establish the cause of its errors. Because we examined this pattern after seeing the aggregate results, the analysis remains exploratory. Reporting both recovery measures alongside local geometry gives a more complete account of the model’s behaviour.

Clustering the full PredNovel population also makes the evaluation relevant to the output a user receives. It includes errors from the known/novel decision and requires no true count of target-private classes. LCC adds a lineage profile after this partition is fixed. Researchers can examine that profile alongside marker expression to guide further work. The Consistent category records whether an evidence-coverage threshold is met. It does not establish cell-type identity or biological correctness. The analysis applies to eligible communities, and its yield varies by dataset. The earlier Zeisel 2018 example illustrates this use, with no independent validation of the primary evaluation. Two further comparisons are needed to assess the method more fully. A deployment comparison should use an external method with the same access to training information and no target class counts. The present strict scBOL analysis shares scoring rules but retains different training and deployment assumptions. A matched experiment that removes or corrupts ontology structure is also needed to separate its contribution from that of the other components. Future work should assess calibration under larger domain shifts and test LCC hypotheses against independent biological evidence. Within these limits, scOLAR provides a practical way to annotate known cells, organise unfamiliar populations and review their possible lineage relationships when the reference is incomplete.

## 4 Conclusions

scOLAR combines annotation of known cell types with detection, clustering and interpretation of unfamiliar populations. It calibrates novelty decisions from reference cells and groups all PredNovel cells without knowing the number of target-private populations in advance. The results support strong novelty ranking, accurate annotation among retained known cells and good recovery when novel types receive equal weight. The method also retains fewer known cells, and its relative performance varies across abundance-weighted and global clustering measures. These trade-offs matter when choosing how many cells to send for further review. The ontology-based descriptions provide hypotheses for that review, with biological identities and lineage assignments requiring independent validation.

## 5 Methods

This section defines the annotation task and cell representation. It then describes the training objectives, inference and lineage interpretation. Figure 1 gives an overview of scOLAR. Algorithm S1 in Additional file 1 specifies the complete procedure.

### 5.1 Problem formulation

The inputs are a labelled reference set 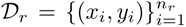 and an unlabelled target set 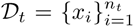. Each cell *x_i_* ∈ R*^G^* is an expression profile over *G* genes, and each reference label *y_i_* belongs to the annotated cell-type set C*_r_*. The target has an unknown cell-type set C*_t_*. The shared types are C*_s_* = C*_r_* ∩ C*_t_*. Novel target types Ĉ*_t_* = C*_t_* \ C*_s_* are absent from the reference. Reference-private types Ĉ*_r_* = C*_r_* \ C*_s_* are absent from the target. The correspondence between C*_r_* and C*_t_* is unknown. The task assigns each target cell either a shared label in C*_s_* or membership in one of the populations discovered within Ĉ*_t_*. The number and identities of novel populations are not supplied.

Training uses both reference and target expression profiles and is therefore transductive. Target labels and the number of target-private classes are withheld from training, calibration, checkpoint selection and inference. Evaluation on D*_t_* takes place after outputs are fixed. In the *intra-dataset* setting, D*_r_* and D*_t_* are cell-disjoint partitions of one dataset, with partly overlapping label sets and a common expression distribution. In the *cross-dataset* setting, they come from separate datasets. Cross-dataset analyses provide historical development context for variation across platforms and protocols.

### 5.2 Cell representations and ontology-indexed prototypes

A shared encoder maps cells to a low-dimensional representation. The classifier compares each cell with learnable representative vectors, called prototypes [48]. Each prototype has a CL term index. Let O = {1*, . . . , N* } index all compiled ontology terms and let K ⊂ O index the terms mapped to observed source classes. This indexing connects the model to the CL relationships used below. The vectors themselves are learned during training and do not come from a pretrained semantic embedding of the ontology.

#### Encoder

The encoder *f*_enc_ : R*^G^* → R*^d^* maps reference and target cells to *d*-dimensional embeddings,

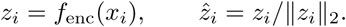

Its input is the standardised matrix of highly variable genes. Raw counts are retained for the generative objective in Section 5.3. Section S1 and Table S3 in Additional file 1 describe the encoder architecture.

#### CL-indexed prototype head

The classifier contains one learnable prototype *µ_j_* ∈ R*^d^* for every compiled term *j* ∈ O,

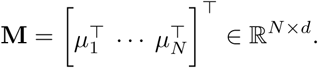

These parameters use Xavier initialisation and share the CL term indices. The K rows correspond to observed source classes and are the only rows eligible for a returned Known label. All *N* rows enter source softmax normalisation and the maximum-logit novelty score. Rows without source examples can therefore compete in scoring. They are not validated zero-shot labels, and their geometry has no established semantic interpretation. Prototypes are *ℓ*_2_-normalised when used in a cosine score.

#### Affinity score

The affinity between cell *i* and prototype *j* is their cosine similarity, multiplied by a single learnable scale *s >* 0. This scale also absorbs the softmax temperature,

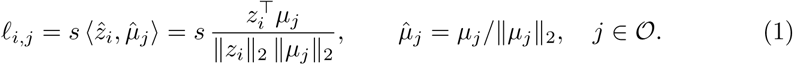

Source classification and MLS use the complete logit vector. The hierarchical centripetal loss (HCL), decision-boundary regularisation (DBR), repulsive term in novel separation (nSep), and post-hoc Lineage Consistency Check (LCC) use the mapped source subset K. The sections below specify which of these components also use CL ancestry or coarse memberships.

### 5.3 Training objectives

Training combines reconstruction, prototype classification, source HCL, source DBR and nSep. CL relationships enter source-label mapping, HCL, DBR and the later LCC analysis. Reconstruction and nSep use no ontology relationships.

#### Generative reconstruction

The model uses a zero-inflated negative binomial (ZINB) likelihood to describe sparse, over-dispersed scRNA-seq counts [49, 50]. The decoder predicts a relative mean *ν̂*, dispersion *θ >* 0 and dropout probability *π* ∈ (0, 1) for each gene. A per-cell size factor *ρ_i_* = *L_i_/*median*_k_*(*L_k_*) rescales the mean to the count range, where *L_i_* = ^Σ^*_g_ x_ig_* is the library size. The size factor is clipped for numerical stability, giving *ν_i_* = *ρ_i_ν_i_*. For an observed count *x*,

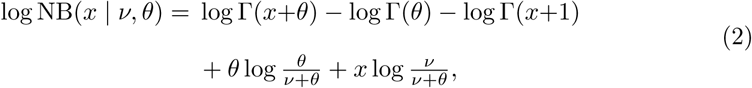

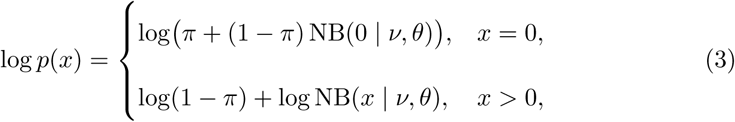

The reconstruction loss L_rec_ = − E [log *p*(*x*)] is averaged over genes and both domains. It trains the representation of unlabelled target cells through their expression profiles.

#### Prototype classification

Reference labels belong to the mapped source subset K. Cross-entropy training normalises the affinity logits in Eq. (1) over the complete prototype head,

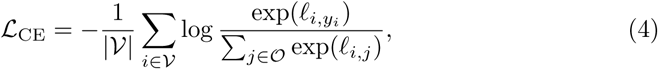

where V indexes reference cells with valid mapped source labels. Mapped rows receive positive supervision. Other rows compete through the normalising denominator. The objective uses no novel target labels.

#### Source hierarchical centripetal objective

HCL uses shared CL ancestors to define positive and negative relationships between mapped source classes. Its margin-based triplet loss follows metric-learning hard mining [51]. The ancestor indicator matrix *A* has *A_ak_*=1 if and only if CL term *k* is an ancestor of mapped source term *a*, including itself. The shared-ancestor count is

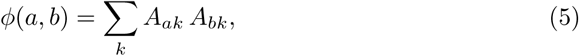

Two mapped source terms are treated as same-lineage when *ϕ*(*a, b*) ≥ *τ* , with *τ* = 3. For an anchor *a*, the positive is the same-lineage candidate with the fewest shared ancestors. The negative is the different-lineage candidate with the most shared ancestors (Section S2 in Additional file 1). For sampled source-class triplets T ,

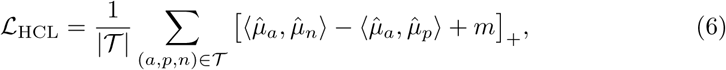

The triplet margin *m* is 0.3 when HCL activates at epoch 20. It decreases to 0.05 at the start of boundary training at epoch 50. The evaluation configuration uses source anchors only and disables target HCL.

#### Decision-boundary regularisation

DBR encourages a correctly classified reference cell to have stronger evidence for its fine type than for a broad lineage. For each coarse ontology term *c* at depth at most *δ* = 3, the model forms a detached coarse reference *µ*^^coarse^. This is the *ℓ*_2_-normalised centroid of the observed prototypes descending from *c*. The strongest lineage affinity of a cell is *S_i_*^coarse^ = max*_c_*⟨*z*^*_i_, µ*^*_c_*^coarse^⟩. For a correctly classified reference cell (*y*^*_i_* = *y_i_*), DBR penalises an insufficient gap between fine-type and lineage affinity. The base margin is *β* = 0.3 and the slack is *η* = 0.05,

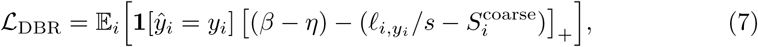

The coarse references are detached CL-derived aggregates, so this term keeps the coarse branch fixed. Gradients act through the fine source-class score and do not propagate through the shared coarse-lineage reference. DBR is applied only to source cells in the evaluation configuration. Target DBR is disabled.

#### Novel separation

A separate term reduces similarity between likely-novel target cells and observed source prototypes. Each target cell receives a soft weight *w_i_* = *σ κ*(*α* − MLS*_i_*) , where *σ* is the logistic sigmoid and *κ* controls its steepness. The weight rises as the score MLS*_i_* over the complete head falls (Section 5.4). The loss is

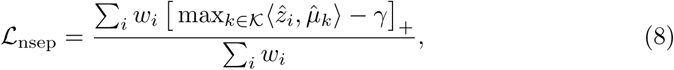

The repulsive maximum uses only the observed source rows K. The small tolerance *γ* allows a narrow band of similarity without penalty. nSep uses no CL ancestry, depth or coarse membership.

#### Overall objective and curriculum

The representation term is L_rep_ = *λ*_rec_L_rec_ + L_CE_, the lineage term is L_lin_ = *λ*_HCL_L_HCL_, and the boundary term is L_bnd_ = *λ*_DBR_L_DBR_ + *λ*_nsep_L_nsep_. Together they give

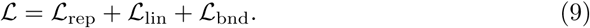

A fixed three-phase schedule activates these terms in sequence. The first phase trains reconstruction and classification (*λ*_rec_ = 1, with the remaining weights zero). The second adds source HCL. The third enables source DBR and nSep and reduces *λ*_rec_. Table S4 in Additional file 1 gives the full schedule and weights.

### 5.4 Open-set inference

Each target cell receives the maximum logit over the complete prototype head, following the maximum-logit criterion for open-set recognition [52–54],

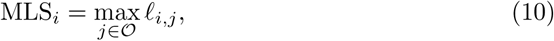

A cell is assigned to PredNovel when MLS*_i_ < α*. Otherwise, its Known label is arg max*_k∈K_ ℓ_i,k_*, restricted to mapped source classes. MLS uses the indexed, learned head and does not directly use ontology relationships.

The threshold *α* is calibrated using reference cells alone. Before training, twenty fixed draws of ten mapped reference classes are sampled. For each draw, cells from the held-out classes are excluded from the calibration score sample. The MLS of every remaining source cell is still calculated over the complete logit set O. The draw-specific threshold is the 5th percentile of these scores, corresponding to a 95% reference-retention target. The final *α* is the median across the twenty draws. Calibration reuses the same draws at epochs 60, 70, 80, 90 and 100. No target expression, label or metric enters this procedure. Neither MLS nor threshold calibration uses CL ancestry or coarse memberships directly. Their connection to the ontology is through the head’s indexing and learned parameters. Section S3 and Algorithm S2 in Additional file 1 give the complete procedure.

For the common-evaluator benchmark, the ground-truth-novel subset and true number of novel classes enter clustering, post-hoc matching and scoring only after predictions and representations are fixed. Deployment uses Leiden [55] at fixed resolution 1.0 on the full PredNovel population, without target labels or true K. LCC reuses this same partition. Strict scBOL also retains the true target class count supplied during its training, even when the downstream clustering step uses no predefined count.

### 5.5 Lineage-consistency interpretation

LCC describes lineage affinities after inference. It reuses the fixed Leiden partition of all PredNovel cells and leaves novelty decisions and clusters unchanged. Source-class affinities are combined through CL ancestry to form a descriptive profile. The profile provides a hypothesis for review and has no independently validated lineage accuracy in this study.

#### Ancestral affinity

For a declared-novel cell, the logits of Eq. (1) are restricted to observed reference types and converted by softmax to a distribution *q_i_*. Let *A*_obs_ be the restriction of the ancestor matrix *A* in Eq. (5) to observed types as rows and all *N* terms as columns.

Accumulating affinity through the ontology gives

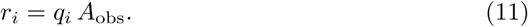

Each fine type contributes to all its ancestors, so the entries of *r_i_* need not sum to one. The vector scores each ontology term by the known-type affinity that flows through it. It therefore ranks lineage affinities, without assigning a fine cell-type identity.

#### Cluster-level judgement

Communities with fewer than five cells are omitted. The remaining communities use the mean of their *r_i_* vectors, with terms shallower than depth 2 discarded. A community is designated *Consistent* if the top three terms reach the predefined 0.40 Top-3 coverage threshold. The deepest of these terms supplies the lineage description. Other communities remain Ambiguous (Section S4 in Additional file 1). The Consistent designation records coverage of the model’s affinity scores. It does not measure biological correctness, validated lineage identity, precision or recall, and it is not a prospectively declared standalone performance endpoint. LCC covers a subset of communities, with substantial variation across datasets.

### 5.6 Datasets and preprocessing

The study uses five intra-dataset benchmarks and three cross-dataset pairs, covering four organisms and five sequencing platforms (Table S1 in Additional file 1). The partitions follow the established benchmark without modification [34]. Shared, reference-private and novel type sets are fixed for each dataset and are not resampled. This class structure is shared by the current scOLAR and strict scBOL comparison. The other methods in Table 1 retain their historical evaluation settings. Cells with annotations outside the benchmark registry are excluded before processing.

All datasets use the same preprocessing procedure. Reference and target expression are joined for unsupervised feature processing. Counts are normalised by library size and log-transformed. Scanpy selects the top 2000 highly variable genes [7]. Each selected gene is standardised over the combined reference–target matrix, and values are clipped to [−10, 10]. The benchmark cell and gene sets are otherwise unchanged. The processed matrix enters the encoder, and raw counts for the same genes supply the ZINB likelihood in Eq. (3). These operations use no target labels.

#### Cao

This whole-organism *Caenorhabditis elegans* dataset uses sci-RNA-seq [40]. It contains 30960 cells across sixteen annotated types, comprising six shared, four reference-private and six novel types. The GEO accession is GSE98561.

#### Quake 10x and Quake Smart-seq2

These are two platform arms of the mouse *Tabula Muris* atlas, covering multiple organs [4]. The 10x arm contains 53587 cells across 36 types (12 shared, 12 reference-private, 12 novel). The Smart-seq2 arm contains 41526 cells across 45 types (15*/*15*/*15). Their shared source and different chemistries motivated the earlier development analyses of platform variation. The GEO accession is GSE109774.

#### Wagner

This zebrafish embryo dataset uses inDrop [41]. It contains 34348 cells across 14 developmental types (5 shared, 4 reference-private, 5 novel). The GEO accession is GSE112294.

#### Zeisel 2018

This mouse nervous system dataset uses 10x [42]. It is the largest benchmark, with 110704 cells across 17 types (6 shared, 5 reference-private, 6 novel). Raw sequence data are available under SRA accession SRP135960.

#### Cross-dataset pairs

Earlier development analyses compare three reference→target pairs across platforms within the same tissue. They comprise mouse mammary gland (Smart-seq2 → 10x, 2405 → 4481 cells), human pancreas (Muraro [56] CEL-Seq2 → Baron [57] inDrop, 1724 → 8451), and human placenta (Smart-seq2 → 10x). Reference and target come from disjoint experiments, so no cell appears in both. These pairs are outside the primary common-evaluator comparison.

### 5.7 Ontology construction

The Cell Ontology release of 2025-12-17 [35] is parsed with obonet and pronto into a directed acyclic graph of *N* = 3238 terms. The release contains 4493 *is-a* edges. The ancestor closure used by HCL, DBR and LCC also follows *develops-from* relations. For each release, preprocessing computes the parent matrix and the ancestor indicator matrix *A* in Eq. (5), both with and without the term itself. It also computes each term’s depth as its distance to the root and a lookup from cell-type names, synonyms and CL identifiers to term indices. The same structures serve all datasets. Dataset annotations are matched through the lookup, without dataset-specific ontology curation.

### 5.8 Training configuration

All datasets within an analysis configuration use the same hyperparameters. Training runs for 100 epochs with AdamW, batch size 1024 and gradient-norm clipping at 5. A fixed cosine schedule reduces the learning rate from 10*^−^*^3^ to 10*^−^*^5^ through all 100 epochs. Validation metrics do not control this schedule. Reference and target loaders are cycled together so that every step includes both domains. For shared classes within an intra-dataset benchmark, each cell enters the reference with probability 0.5 and otherwise enters the target. Reference-private cells remain in the reference, and target-private cells remain in the target. Every reference cell used for source training has its source label.

In the primary evaluation, target loaders contain expression only. They exclude all target ground-truth fields. Target labels enter scoring after predictions are fixed and are excluded from optimisation, checkpoint selection, reference-only held-out-class calibration, inference, clustering and LCC. Target HCL, target DBR and target-neighbourhood consistency (TNC) are disabled.

The three phases in Eq. (9) follow a fixed schedule. Phase P1 (epochs 1–19) trains L_rec_ +L_CE_ alone. Phase P2 (epochs 20–49) adds HCL at *λ*_HCL_ = 0.1. Its triplet margin decreases linearly from 0.3 to 0.05 across the phase. The evaluation uses *λ*_HCL_ = 0.1 on the source side and disables target HCL. Unlabelled target cells contribute to reconstruction and nSep, with target DBR and TNC disabled.

Phase P3 (epochs 50–100) activates DBR at *λ*_DBR_ = 0.05, with base margin *β* = 0.3 and slack *η* = 0.05. The reconstruction weight falls from 1.0 to 0.5 at this transition, then decreases linearly to zero between epochs 80 and 100. Threshold calibration occurs at epochs 60, 70, 80, 90 and 100. From epoch 60, nSep is active at *λ*_nsep_ = 0.01, with steepness *κ* = 5.0 and tolerance *γ* = 0.1. The same-lineage threshold is *τ* = 3 shared ancestors, and the coarse-term depth bound is *δ* = 3.

Every scOLAR run completes the 100-epoch schedule and uses the final epoch-100 checkpoint fixed before target scoring. Target labels select no epoch, checkpoint, threshold, clustering resolution or model. They enter only the predefined benchmark evaluations after outputs are fixed. Source HCL and source DBR follow the schedule, with target HCL, target DBR and TNC off. Table S4 in Additional file 1 lists the complete configuration.

### 5.9 Evaluation metrics

Summaries use four independent runs. The common-evaluator comparison reports Known classification on ground-truth-known cells and Novel weighted, Novel macro, ARI and OverallJ on the predefined benchmark outputs. The clustering evaluation uses ground-truth-novel membership and true *K* only after predictions and representations are fixed. Cluster labels are matched post hoc with the Hungarian algorithm [58].

Deployment evaluates the model’s PredNovel decisions and the fixed Leiden partition of all PredNovel cells. Target ground truth and true K are excluded from these decisions. We report known retention, accuracy conditional on retention and the joint correct-known rate separately. The joint rate is calculated for individual cells, so it cannot be recovered by multiplying aggregate retention and conditional accuracy. AUROC and average precision assess score ranking independently of a chosen threshold. Additional file 1 explains the score used for each method.

### 5.10 Implementation and runtime

scOLAR uses PyTorch 2.8, Python 3.9 and CUDA 12.8. Preprocessing uses Scanpy 1.10.3 and AnnData 0.10.8. Ontology parsing uses obonet 1.1.1 and pronto 2.7.3. The recorded development runtimes cover preprocessing, ontology construction, training, threshold calibration, inference and lineage analysis on NVIDIA RTX 3090 and RTX 4090 GPUs (Table S8 in Additional file 1). Replicate identities, the definition of the recorded spread and the dataset-to-GPU assignments are unavailable in those records. These times provide computational context and cannot support a hardware-normalised comparison.

### 5.11 Use of large language models

OpenAI ChatGPT and Codex assisted with language editing, figure scripting and formatting, code review, and consistency checks. The authors reviewed and verified all scientific analyses, numerical results, interpretations, citations and final manuscript content.

### 5.12 Baseline comparison

Strict scBOL uses the same evaluation populations and scoring rules as scOLAR. Its training receives the true target class count, and its checkpoint is fixed before target scoring. The shared evaluator therefore preserves differences in training information and deployment assumptions. Clustering ground-truth-novel cells with true *K* is a benchmark-only step. The true target class count supplied to scBOL during training remains an additional input.

The published scNym, scArches, ItClust, MARS, scCNC and scDECL results, together with earlier scBOL/scOLAR analyses, provide historical context. Their training information, selection rules and deployment assumptions differ and were not standardised in this study (Table S2 in Additional file 1).

## Abbreviations

AUROC: area under the receiver operating characteristic curve
CE: cross-entropy
CL: Cell Ontology
DBR: decision-boundary regularisation
HCL: hierarchical centripetal loss
LCC: Lineage Consistency Check
MLS: maximum logit score
NB: negative binomial
scRNA-seq: single-cell RNA sequencing
UMAP: uniform manifold approximation and projection
ZINB: zero-inflated negative binomial

## Additional files

**Additional file 1.** *File format.* PDF (.pdf). *Title of data.* Supplementary Methods and Results for scOLAR. *Description of data.* Complete results under the common evaluator, deployment performance and end-to-end known-cell summaries, the exploratory Quake Smart-seq2 analysis, descriptive LCC results, implementation details and dataset sources. Earlier development analyses are identified separately.

## Declarations

### Ethics approval and consent to participate

Not applicable.

### Consent for publication

Not applicable.

### Availability of data and materials

All datasets supporting the conclusions of this article are publicly available published benchmarks. The standardised open-set partitions follow the protocol of Zhai et al. [34]. Table S1 in Additional file 1 lists verified accession identifiers for the five intra-dataset benchmarks and the public source studies for the three cross-dataset pairs. The scOLAR version 1.0.0 source code is publicly available from GitHub, and the exact release is archived on Zenodo under the MIT license. Section S6 and Table S8 in Additional file 1 describe the recorded software, hardware and development runtimes.

^ Project name: scOLAR

^ Project home page: https://github.com/Rijinna/scOLAR-project

^ Archived version: https://doi.org/10.5281/zenodo.22262387

^ Operating system(s): Platform independent

^ Programming language: Python

^ Other requirements: Python 3.9 or higher, PyTorch 2.8

^ License: MIT

^ Any restrictions to use by non-academics: None

## Authors’ contributions

YL designed the method, implemented the experiments, analyzed the results, and wrote the manuscript. SY contributed to methodological discussions and manuscript revision. HY contributed to experimental discussions and manuscript revision. WJ supervised the project, secured funding, and revised the manuscript. All authors read and approved the final manuscript.

## Acknowledgements

Not applicable.

