## Additional File 1 for "scOLAR: Ontology-Anchored Open-Set Annotation of Single-Cell RNA-seq Data"

This supplement describes the datasets, implementation, training and evaluation of scOLAR. It includes results that support the main text and identifies earlier development analyses separately. The primary analysis uses four independent runs and the final epoch-100 checkpoint. Target ground truth enters benchmark scoring after predictions are fixed. It is excluded from training, inference, checkpoint selection, reference-only held-out-class calibration, clustering and LCC.

### S1 Additional details of datasets, model and baselines

#### S1.1 Datasets and open-set partitions

The five primary scRNA-seq benchmarks are Cao [1], Quake_10x and Quake_Smart-seq2 [2], Wagner [3] and Zeisel_2018 [4]. Earlier development analyses also use cross-dataset pairs from the same organ on different platforms. These cover mammary gland [2], pancreas (Muraro [19] → Baron [20]) and placenta (Vento-Tormo [18]). The open-set partitions follow the published benchmark without modification [5]. The shared ($C_s$), reference-private ($\bar{C}_r$) and target-private or novel ($\bar{C}_t$) type sets are fixed for each dataset.

Reference and target contain disjoint cells and partly overlapping label sets. For shared classes, each cell enters the reference with probability 0.5 and otherwise enters the target. Reference-private cells remain in the reference, and target-private cells remain in the target. Every cell used for source training has its reference label. Table S1 lists the datasets, cell counts, public sources and type partitions. Published baseline values retain their original evaluation settings.

**Table S1.** Datasets, public sources and open-set partitions used in this study. $|C_s|$, $|\bar{C}_r|$ and $|\bar{C}_t|$ are the numbers of shared, reference-private and novel (target-private) cell types.

| Dataset | Public source | Organism / tissue · platform | Cells | $\lvert C_s\rvert$ | $\lvert\bar{C}_r\rvert$ | $\lvert\bar{C}_t\rvert$ |
|---|---|---|---|---|---|---|
| *Intra-dataset benchmarks* | | | | | | |
| Cao | GEO GSE98561 | *C. elegans* · sci-RNA-seq | 30 960 | 6 | 4 | 6 |
| Quake_10x | GEO GSE109774 | Mouse atlas · 10x | 53 587 | 12 | 12 | 12 |
| Quake_ Smart-seq2 | GEO GSE109774 | Mouse atlas · Smart-seq2 | 41 526 | 15 | 15 | 15 |
| Wagner | GEO GSE112294 | Zebrafish embryo · inDrop | 34 348 | 5 | 4 | 5 |
| Zeisel_2018 | SRA SRP135960 | Mouse nervous system · 10x | 110 704 | 6 | 5 | 6 |
| *Cross-dataset pairs (reference → target)* | | | | | | |
| Mammary gland | Tabula Muris [2]; GEO GSE109774 | Mouse · SS2 → 10x | 2 405 / 4 481 | 4 | 0 | 3 |
| Pancreas | Muraro [19] → Baron [20] | Human · CEL-Seq2 → inDrop | 1 724 / 8 451 | 4 | 0 | 4 |
| Placenta | Vento-Tormo [18] | Human · SS2 → 10x | 4 310 / 54 976 | 4 | 0 | 4 |

*Note.* Cell counts and type partitions follow the benchmark of [5] (its Tables S2–S4). Verified accessions are given where they are recorded in the manuscript sources; otherwise the public source study is identified. For cross-dataset pairs, the two figures are reference and target cell counts.

#### S1.2 Preprocessing

Scanpy [6] preprocesses reference and target cells together. All statistics are computed on $D_r \cup D_t$, so target cells require no separate normalisation at inference. Raw integer counts come from `adata.layers['counts']`, or from `adata.X` if that layer is absent. The counts remain unchanged for the ZINB likelihood. The encoder input is prepared as follows.

1. Library sizes are normalised to $10^4$ counts per cell with `sc.pp.normalize_total`.
2. Values are log-transformed as $x \mapsto \log(1 + x)$ with `sc.pp.log1p`.
3. The $G = 2000$ most highly variable genes are selected with `sc.pp.highly_variable_genes`. The call sets `n_top_genes` to 2000 and leaves `subset` false. It returns a Boolean mask, which is applied to obtain the input matrix.
4. Each gene is standardised to zero mean and unit variance. Values are clipped to $[-10, 10]$ to limit extreme $z$-scores.

The benchmark cell and gene sets are already quality-controlled and remain unchanged. The ZINB size factor is $\rho_i = L_i/\mathrm{median}_k(L_k)$, where $L_i = \sum_g x_{ig}$ is the raw library size. It is clipped to $[0.1, 10]$ for numerical stability.

#### S1.3 Cell Ontology mapping and prototype-head indexing

The Cell Ontology (CL) [7] release `2025-12-17` is parsed with `obonet` 1.1.1 and compiled into one static graph for all runs. Obsolete entries and terms outside the `is_a` and `develops_from` closure of `CL:0000000` are removed. The resulting graph contains $N = 3238$ terms. The release has 4493 `is_a` edges. This count excludes the additional relation type used in the ancestor closure. The compiled term index also defines the $N$ prototype-head rows. Source labels are mapped to CL terms, with $\mathcal{K}$ indexing the mapped subset. The implementation raises an error if a benchmark source cell has no usable mapping. Target-private labels are excluded from training.

The ancestor indicator matrix $A \in \{0,1\}^{N\times N}$ follows the transitive closure of `is_a` and `develops_from` edges. An entry $A_{ak} = 1$ means that term $k$ is an ancestor of term $a$, including itself. The full graph supplies source HCL relationships, coarse memberships for source DBR, and ancestry for post-hoc LCC. The parameter array $M_{\text{impl}} \in \mathbb{R}^{N\times d}$ contains one Xavier-uniform row per compiled term. Source cross-entropy and MLS use all $N$ rows. Known labels can only come from the mapped subset $\mathcal{K}$. Source HCL, source DBR, the nSep repulsive similarity and LCC also use that subset. Rows without source examples remain competing classifier parameters. They are not pretrained or validated zero-shot labels, and their geometry has no established semantic interpretation.

#### S1.4 Baseline implementations

Published MARS, ItClust, scNym, scArches, scCNC and scDECL results, together with earlier scBOL/scOLAR analyses, provide historical context. Their evaluators, training information and deployment assumptions differ and were not standardised in this study. Strict scBOL is evaluated separately with the common evaluator and retains the true target class count supplied during training. Table S2 describes the historical method families.

**Table S2.** Seven baseline methods in the historical benchmark. Under that benchmark protocol, methods in the first family return a known label or a generic *unassigned* label, and $k$-means groups the unassigned cells. The second family clusters cells without identifying known types. Among the listed baselines, scBOL combines known-cell annotation with novel-cell clustering. The historical rows use different evaluation settings.

| Method | Year | Family | Novel populations obtained by | Implementation |
|---|---|---|---|---|
| MARS [8] | 2020 | Known + unassigned | $k$-means on unassigned cells | `github.com/snap-stanford/mars` |
| ItClust [9] | 2020 | Known + unassigned | $k$-means on unassigned cells | `github.com/jianhuupenn/ItClust` |
| scNym [10] | 2021 | Known + unassigned | $k$-means on unassigned cells | `github.com/calico/scnym` |
| scArches [11] | 2022 | Known + unassigned | $k$-means on unassigned cells | `github.com/theislab/scarches` |
| scCNC [12] | 2022 | Unsupervised clustering | Clustering output | `github.com/WHY-17/scCNC` |
| scDECL [13] | 2023 | Unsupervised clustering | Clustering output | `github.com/DBLABDHU/scDECL` |
| scBOL [5] | 2024 | Joint known/novel | Bipartite prototype alignment | `github.com/aimeeyaoyao/scBOL` |
| scOLAR (ours) | — | Joint known/novel | Leiden on declared-novel cells | `github.com/Rijinna/scOLAR-project` |

*Note.* Historical rows retain their original evaluators. The common benchmark uses ground-truth-novel membership and true $K$ for clustering only after outputs are fixed. Deployment groups each method's PredNovel cells with fixed Leiden, without target labels or true K. Strict scBOL still receives the true target class count during training.

#### S1.5 Network architecture

The encoder $f_{\text{enc}} : \mathbb{R}^G \to \mathbb{R}^d$ is a three-layer perceptron with layer normalisation and LeakyReLU activations. It maps a standardised 2000-dimensional highly-variable-gene (HVG) vector to a latent code of dimension $d = 128$. The symmetric decoder has three heads for the ZINB parameters. A softmax over genes gives the relative mean $\bar{\nu}$, softplus gives dispersion $\theta > 0$, and sigmoid gives dropout probability $\pi \in (0,1)$.

The prototype array is stored by rows as $M_{\text{impl}} = M \in \mathbb{R}^{N\times d}$, matching the main-text notation. This array and the learnable logit scale $s > 0$, initialised at 8.0, are the only parameters outside the encoder–decoder path. All objectives use the shared cell representation. Source HCL and source DBR additionally use CL ancestry or coarse memberships. Table S3 gives the same layer configuration used for every dataset.

### S2 Optimisation and training strategy

The following symbols are used in the main text and this supplement.

**Table S3.** Architecture of scOLAR. LN denotes layer normalisation. Activations are LeakyReLU with negative slope 0.2.

| Module | Layer | Operation | Output dim. |
|---|---|---|---|
| Encoder | Input | Standardised HVG matrix | 2000 |
| | Hidden 1 | Linear + LN + LeakyReLU + Dropout(0.2) | 512 |
| | Hidden 2 | Linear + LN + LeakyReLU | 256 |
| | Latent | Linear + LN | $d = 128$ |
| Decoder | Hidden 1 | Linear + LeakyReLU, then Linear + LeakyReLU | $256 \to 512$ |
| | Mean head $\bar{\nu}$ | Linear + softmax over genes | 2000 |
| | Dispersion head $\theta$ | Linear + softplus | 2000 |
| | Dropout head $\pi$ | Linear + sigmoid | 2000 |
| Prototype head | CL-indexed prototypes $M_{\text{impl}} = M$ | Xavier-uniform, $\ell_2$-normalised in Eq. (1) | $N \times 128$ |
| | Logit scale $s$ | Learnable, initialised at 8.0 | 1 |

*Note.* The latent code $z_i$ enters both the decoder and the cosine affinity in main-text Eq. (1). Source DBR uses detached coarse references formed from CL relationships. The coarse branch remains fixed for this term. Gradients act through the fine source-class score and do not propagate through the shared coarse-lineage reference (Section S3.2). Mini-batches with non-finite activations are detected and skipped.

| Symbol | Meaning | Symbol | Meaning |
|---|---|---|---|
| $D_r, D_t$ | Reference and target set | $\ell_{i,j}$ | Affinity of cell $i$ to ontology-indexed prototype $j$, Eq. (1) |
| $C_s$ | Shared cell types | $s$ | Learnable logit scale |
| $\bar{C}_r$ | Reference-private types | $\text{MLS}_i$ | Maximum full-head logit, Eq. (10) |
| $\bar{C}_t$ | Novel (target-private) types | $\alpha$ | Novelty threshold |
| $G$ | Number of input genes | $\phi(a,b)$ | Shared-ancestor count, Eq. (5) |
| $d$ | Latent dimension | $\tau$ | Same-lineage threshold on $\phi$ |
| $N$ | Number of CL graph terms | $\delta$ | Depth bound for coarse terms |
| $M$ | Learned CL-indexed $N \times d$ prototype head | $\beta, \eta$ | DBR base margin and slack, Eq. (7) |
| $O$ | All compiled CL term indices | $\kappa, \gamma$ | Novelty steepness and tolerance, Eq. (8) |
| $\mathcal{K}$ | Mapped observed source subset | $\mathcal{V}$ | Mapped reference cells in Eq. (4) |
| $A$ | Ontology ancestor matrix | $m$ | Triplet margin, Eq. (6) |
| $z_i, \hat{z}_i$ | Latent code and its unit norm | $q_i, r_i$ | Known-type and lineage affinity, Eq. (11) |
| $\mu_j, \hat{\mu}_j$ | Ontology-indexed prototype and its unit norm | $\rho_i$ | Per-cell size factor |

### S2.1 Triplet construction for the hierarchical centripetal objective

The hierarchical centripetal loss (HCL) in main-text Eq. (6) uses CL relationships to guide the geometry of source-class prototypes. Each triplet $(a, p, n)$ contains mapped source rows only. Unobserved ontology terms do not serve as pseudo-labelled source classes. The margin follows a metric-learning triplet loss [14], with positive and negative candidates defined by CL ancestry. Two source terms are treated as same-lineage when their shared-ancestor count $\phi(a,b) = \sum_k A_{ak}A_{bk}$ reaches $\tau = 3$. Deeper terms have more ancestors, so $\phi$ is used only for this binary split. It is not interpreted as a graded distance.

Anchors are mapped source terms present in the current source mini-batch. For anchor $a$, the candidate pools are $P(a) = \{b : \phi(a,b) \geq \tau,\ b \neq a\}$ and $N(a) = \{b : \phi(a,b) < \tau\}$. Hard mining selects the positive in $P(a)$ with the fewest shared ancestors and the negative in $N(a)$ with the most. Ties are sampled without using prototype cosine similarity. Triplets are resampled for each batch. The margin decreases linearly from $m_{\text{start}} = 0.3$ when HCL activates to $m_{\text{end}} = 0.05$ at the start of the boundary phase. It remains at 0.05 thereafter, allowing finer adjustment after the initial lineage separation. The shared-ancestor threshold defines how triplets are constructed and does not establish biological correctness.

### S2.2 Source-only HCL

The evaluation configuration constructs hierarchical triplets from source anchors only. Target HCL, target DBR and target-neighbourhood consistency (TNC) are disabled. Unlabelled target cells contribute to reconstruction and novel separation (nSep).

### S2.3 Curriculum and hyperparameters

Training runs for 100 epochs in three fixed phases. P1 (epochs 1–19) uses reconstruction and source-class cross-entropy. P2 (epochs 20–49) activates source HCL and reduces its triplet margin. P3 (epochs 50–100) activates source decision-boundary regularisation (DBR). Reference-only held-out-class calibration occurs at epochs 60, 70, 80, 90 and 100, with nSep active from epoch 60. Table S4 gives the full schedule and hyperparameters.

At epoch $t \in \{1, \dots, 100\}$, AdamW uses the fixed cosine learning-rate schedule

$$\mathrm{lr}_t = 10^{-5} + \tfrac{1}{2}(10^{-3} - 10^{-5}) \left[1 + \cos\!\left(\pi \frac{t-1}{99}\right)\right].$$

The learning rate starts at $10^{-3}$ and reaches $10^{-5}$ at epoch 100. Validation metrics do not control the schedule.

**Table S4.** Training schedule and hyperparameters of scOLAR. All values are shared across datasets. Symbols follow the main text.

| Group | Item | Value | Note |
|---|---|---|---|
| *Curriculum (100 epochs total)* | | | |
| | P1 warm-up | epochs 1–19 | $\mathcal{L}_{\text{rec}} + \mathcal{L}_{\text{CE}}$ only |
| | P2 source HCL | epochs 20–49 | $+\,\mathcal{L}_{\text{HCL}}$; margin 0.3 at activation, 0.05 at epoch 50 |
| | P3 boundary refinement | epochs 50–100 | $+\,\mathcal{L}_{\text{DBR}}$; $\lambda_{\text{rec}}$ halved |
| | calibration grid | epochs 60, 70, 80, 90, 100 | fixed reference-class draws, $+\,\mathcal{L}_{\text{nsep}}$ from epoch 60 |
| | reconstruction ramp | epochs 80–100 | $\lambda_{\text{rec}}$ decays linearly $0.5 \to 0$ |
| *Loss weights* | | | |
| | Prototype classification | 1 (fixed) | Main-text Eq. (9) |
| | Reconstruction $\lambda_{\text{rec}}$ | $1.0 \to 0.5 \to 0$ | 1.0 (ep. 1–49), 0.5 (ep. 50–79), linear to 0 by ep. 100 |
| | Hierarchical $\lambda_{\text{HCL}}$ | 0.1 | Source side only |
| | target-side factor (primary) | 0 | Section S2.2 |
| | Decision boundary $\lambda_{\text{DBR}}$ | 0.05 | Active from epoch 50 |
| | Novel separation $\lambda_{\text{nsep}}$ | 0.01 | Active from epoch 60 |
| *Objective-specific* | | | |
| | Same-lineage threshold $\tau$ | 3 | Shared CL ancestors |
| | Triplet margin $m$ | $0.3 \to 0.05$ | Linearly annealed over P2 |
| | Coarse-term depth bound $\delta$ | 3 | Main-text Eq. (7) |
| | DBR margin $\beta$ | 0.3 | Required fine-minus-coarse cosine gap |
| | DBR slack $\eta$ | 0.05 | Hinge on $(\beta - \eta) - \text{gap}$ |
| | Novelty-weight steepness $\kappa$ | 5.0 | Main-text Eq. (8) |
| | Separation tolerance $\gamma$ | 0.1 | Main-text Eq. (8) |
| *Optimisation* | | | |
| | Optimiser | AdamW | weight decay $10^{-4}$ |
| | Learning rate | $10^{-3} \to 10^{-5}$ | Fixed cosine schedule over all 100 epochs |
| | Batch size | 1024 | Reference and target loaders cycled jointly |
| | Reference allocation | 0.5 | Probability that a shared-class cell enters the reference |
| | Gradient clipping | $\lVert g \rVert_2 \le 5$ | Global norm |
| | Checkpoint selection | fixed final epoch 100 | Fixed before target scoring, without target labels |
| *Inference* | | | |
| | Known TPR at calibration | 95% ($\epsilon = 0.05$) | Section S3 |
| | LCC softmax temperature | 2.0 | Section S4.1 |
| | Novel clustering | Leiden [17], resolution 1.0 | Section S4.2 |

*Note.* All primary runs complete the 100-epoch curriculum and use the epoch-100 checkpoint fixed before target scoring. Target ground truth does not enter optimization, inference, checkpoint selection, reference-only held-out-class calibration, clustering or LCC.

#### S2.4 Complete algorithm

Algorithm S1 gives the complete training and inference procedure outlined in main-text Section 2.

### S3 Open-set calibration and boundary-shaping objectives

#### S3.1 Reference-only held-out-class calibration

The maximum logit score (MLS) is $\mathrm{MLS}_i = \max_{j \in O} \ell_{i,j}$ over the complete head [15,16]. Comparing this score with $\alpha$ gives the known/novel decision. MLS and threshold calibration use the head's indices and learned parameters. They do not directly use CL ancestry, depth or coarse memberships.

Before training, twenty draws of ten mapped reference classes are fixed. For each draw, cells from the held-out classes are excluded from the calibration score sample. The MLS of every remaining source cell still uses all $j \in O$. The threshold for each draw is the $\epsilon$-quantile with $\epsilon = 0.05$, targeting 95% reference retention. The final threshold is the median across the twenty draws (Algorithm S2). Calibration reuses those draws at epochs $60, 70, 80, 90$ and 100. No target cells, labels or metrics enter this procedure.

**Algorithm S1** scOLAR training, open-set inference and lineage interpretation.

**Require:** Labelled reference $D_r = \{(x_i, y_i)\}$, unlabelled target $D_t = \{x_i\}$, Cell Ontology ancestor matrix $A \in \{0,1\}^{N\times N}$, phase boundaries $T_1 = 20$, $T_2 = 50$, $T_3 = 60$, calibration grid $C_{\text{cal}} = \{60, 70, 80, 90, 100\}$, total epochs $T = 100$, hyperparameters of Table S4
**Ensure:** Known-class labels for target cells, novel populations, and descriptive post-hoc lineage profiles

Let $O = \{1, \ldots, N\}$ index compiled CL terms; map source labels to the subset $\mathcal{K} \subset O$ ▷ Section S1.3
Initialise encoder, ZINB decoder, $N \times d$ prototype head $M$ and logit scale $s$; register the ancestor and fine-to-coarse buffers
Draw and freeze $R = 20$ held-out reference-class sets $H_r \subset \mathcal{K}$
**for** $t = 1$ to $T$ **do**
  Set the learning rate by the fixed cosine schedule
  Set $(\lambda_{\text{rec}}, \lambda_{\text{HCL}}, \lambda_{\text{DBR}}, \lambda_{\text{nsep}})$ and the triplet margin from the phase schedule
  **if** $t \in C_{\text{cal}}$ **then**
    $\alpha \leftarrow$ CalibrateThreshold$(D_r, O, \mathcal{K}, \{H_r\}_{r=1}^{R})$ ▷ Algorithm S2; reference only
  **end if**
  **for** each paired mini-batch $(B_r, B_t)$ **do**
    $z_i \leftarrow f_{\text{enc}}(x_i)$; $\hat{z}_i \leftarrow z_i/\|z_i\|_2$; $\ell_{i,j} \leftarrow s\,\langle \hat{z}_i, \hat{\mu}_j\rangle$ for $j \in O$ ▷ Eq. (1), full prototype head
    $\mathcal{L}_{\text{rec}} \leftarrow$ ZINB negative log-likelihood on raw counts, over $B_r$ *and* $B_t$
    $\mathcal{L}_{\text{CE}} \leftarrow$ cross-entropy over $j \in O$ on reference cells with a valid CL mapping
    **if** $t \geq T_1$ **then**
      Sample mapped source-class triplets $(a, p, n)$ with CL-based hard mining ▷ Section S2.1
      $\mathcal{L}_{\text{HCL}} \leftarrow$ margin ranking loss on ancestor-mined reference triplets ▷ primary configuration
    **end if**
    **if** $t \geq T_2$ **then**
      $S_i^{\text{coarse}} \leftarrow \max_{c:\text{depth}(c)\leq\delta} \langle \hat{z}_i, \hat{\mu}_c^{\text{coarse}}\rangle$, with $\hat{\mu}^{\text{coarse}}$ detached
      $\mathcal{L}_{\text{DBR}} \leftarrow$ hinge on $(\beta - \eta) - (\ell_{i,y_i}/s - S_i^{\text{coarse}})$ over correctly classified reference cells
    **end if**
    **if** $t \geq T_3$ **then**
      $w_i \leftarrow \sigma(\kappa(\alpha - \text{MLS}_i))$ on target cells; $\mathcal{L}_{\text{nsep}} \leftarrow$ weighted hinge of Eq. (8)
    **end if**
    $\mathcal{L} \leftarrow \lambda_{\text{rec}}\mathcal{L}_{\text{rec}} + \mathcal{L}_{\text{CE}} + \lambda_{\text{HCL}}\mathcal{L}_{\text{HCL}} + \lambda_{\text{DBR}}\mathcal{L}_{\text{DBR}} + \lambda_{\text{nsep}}\mathcal{L}_{\text{nsep}}$
    Clip $\|\nabla\mathcal{L}\|_2$ to 5 and update all parameters with AdamW
  **end for**
**end for**
At epoch $T = 100$, save the final checkpoint before target scoring
Use that final checkpoint for all target inference and evaluation
$\text{MLS}_i \leftarrow \max_{j\in O} \ell_{i,j}$ for every $x_i \in D_t$
$\mathcal{N} \leftarrow \{i : \text{MLS}_i < \alpha\}$; assign $\arg\max_{k\in\mathcal{K}} \ell_{i,k}$ to every $i \notin \mathcal{N}$
Cluster $\{\hat{z}_i\}_{i\in\mathcal{N}}$ with Leiden into candidate novel populations
**for all** novel clusters $C$ **do**
  Omit $C$ from LCC if $|C| < 5$
  $r_i \leftarrow q_i A_{\text{obs}}$ for $i \in C$; average over $C$ and discard terms of depth $< 2$ ▷ Eq. (11)
  Report *Consistent* and the deepest top-three context if coverage is $\geq 40\%$; otherwise report *Ambiguous*
**end for**
**return** known labels, novel populations, descriptive lineage profiles

**Algorithm S2** `CalibrateThreshold` estimates the novelty threshold $\alpha$ from reference cells using fixed held-out-class draws.

**Require:** Reference set $D_r$, full term-index set $O$, mapped source-class subset $\mathcal{K}$, fixed held-out-class draws $\{H_r\}_{r=1}^{R}$ with $|H_r| = 10$, $R = 20$, tolerance $\epsilon = 0.05$
**Ensure:** Threshold $\alpha$

**for** $r = 1$ to $R$ **do**
  Reuse fixed $H_r \subset \mathcal{K}$ ▷ held-out reference classes
  $\mathcal{S}_r \leftarrow \{\max_{j\in O} \ell_{i,j} : x_i \in D_r,\ y_i \notin H_r\}$
  $\alpha_r \leftarrow \text{quantile}(\mathcal{S}_r, \epsilon)$
**end for**
**return** $\alpha \leftarrow \text{median}(\alpha_1, \ldots, \alpha_R)$

The operating point is therefore selected without optimisation against the target dataset. Earlier cross-dataset threshold analyses provide development context only. They do not establish a general calibration or transfer mechanism.

#### S3.2 Decision-boundary regularisation

For each coarse ontology term $c$ at depth at most $\delta = 3$, $\hat{\mu}_c^{\mathrm{coarse}}$ is the $\ell_2$-normalised, equally weighted centroid of the observed source prototypes descending from $c$. CL relationships determine which prototypes enter each centroid. These coarse references are detached, keeping the coarse branch fixed for this term. Gradients act through the fine source-class score and do not propagate through the shared coarse-lineage reference. DBR penalises a fine-minus-coarse cosine gap below $\beta - \eta$, using base margin $\beta = 0.3$ and slack $\eta = 0.05$. The loss applies only to correctly classified source cells ($\hat{y}_i = y_i$). Target DBR is disabled in the evaluation configuration.

#### S3.3 Novel separation

The novelty weight $w_i = \sigma(\kappa(\alpha - \mathrm{MLS}_i))$ with $\kappa = 5.0$ uses MLS over the complete head. It provides a soft novelty weight during training, without a hard known/novel assignment. Weights are detached, and the loss is normalised by $\sum_i w_i$. This keeps its scale from depending on how many target cells currently appear novel. Without normalisation, the term can become negligible when there are few novel cells and dominate when there are many. nSep starts at epoch 60, after the first calibration provides a finite $\alpha$. Its repulsive similarity uses only mapped source rows $\mathcal{K}$. The term uses no CL ancestry, depth or coarse memberships.

### S4 Lineage-consistency interpretation

#### S4.1 Ancestral affinity

The Lineage Consistency Check (LCC) reuses the fixed Leiden partition of all PredNovel cells. For each PredNovel cell, the source-class logits in main-text Eq. (1) are converted to $q_i$ by softmax with temperature $T = 2.0$. Affinity is accumulated through CL ancestry as $r_i = q_i A_{\mathrm{obs}}$. The rows of $A_{\mathrm{obs}}$ index mapped source classes, and its columns index all $N$ graph terms. The resulting vector describes lineage affinity after clustering. It does not establish a ground-truth cell type or a validated lineage assignment.

#### S4.2 Cluster-level judgement

Leiden [17] partitions PredNovel cells once at fixed resolution 1.0, without target labels or true K. LCC keeps this partition unchanged. Communities with fewer than five cells are omitted. For each remaining community, the vectors $r_i$ are averaged to $\bar{r}_C$, and terms at depth $< 2$ are discarded. Let $\tilde{r}_C$ be this depth-filtered vector and $t_{(1)}, t_{(2)}, t_{(3)}$ its three highest-scoring terms. A community meets the *lineage-consistent* criterion when

$$\frac{\tilde{r}_C[t_{(1)}] + \tilde{r}_C[t_{(2)}] + \tilde{r}_C[t_{(3)}]}{\sum_t \tilde{r}_C[t]} \geq 0.40, \tag{S1}$$

It then receives the designation *Consistent*. Other communities receive *Ambiguous*. The deepest of the top three terms supplies the lineage description for a Consistent community. This designation records whether the predefined Top-3 coverage threshold is met. It does not measure biological precision, validated correctness or independently verified lineage identity, and it is not a standalone performance endpoint. LCC covers a subset of communities, and its yield varies substantially across datasets.

### S5 Common-evaluator and deployment evaluation

#### S5.1 Common-evaluator benchmark

The common evaluator tests how scOLAR and strict scBOL perform when outputs are scored on the same benchmark populations. Ground-truth-novel membership and true $K$ enter benchmark clustering only after predictions and representations are fixed. Strict scBOL also receives the true target class count during training. The comparison therefore shares scoring rules while preserving each method's training information and deployment assumptions.

**Table S5. Common-evaluator benchmark.** Values are means ± sample SDs across four independent runs. Every row has $n = 4$. Paired deltas are scOLAR minus strict scBOL.

| Metric | scOLAR | strict scBOL | Paired delta (scOLAR minus scBOL) |
|---|---|---|---|
| Known | 0.9682 ± 0.0006 | 0.9584 ± 0.0064 | 0.0098 ± 0.0065 |
| Novel weighted | 0.6389 ± 0.0225 | 0.6656 ± 0.0319 | −0.0267 ± 0.0531 |
| Novel macro | 0.6586 ± 0.0187 | 0.5867 ± 0.0197 | 0.0720 ± 0.0367 |
| ARI | 0.5093 ± 0.0277 | 0.5280 ± 0.0382 | −0.0188 ± 0.0623 |
| OverallJ | 0.7549 ± 0.0143 | 0.7673 ± 0.0232 | −0.0124 ± 0.0368 |

*Note.* Ground-truth-novel cells and true $K$ are used for benchmark clustering after outputs are fixed. This analysis evaluates the complete systems under benchmark conditions. It does not measure native deployment or isolate the contribution of ontology structure. Strict scBOL receives the true target class count during training.

All four strict-scBOL runs remain in the summary. One valid Zeisel_2018 run has Known 0.985009, Novel weighted 0.470293, Novel macro 0.557001, ARI 0.266257 and OverallJ 0.617958. Its poorer novel clustering is consistent with unstable cluster allocation, although the analysis identifies no operational failure or definitive mechanism in the learned representation.

#### S5.2 Native deployment

Deployment begins with each method's own PredNovel decisions. scOLAR compares MLS with its reference-calibrated threshold, then partitions all PredNovel cells with fixed Leiden. These decisions use no target labels or true K. Strict scBOL's deployment results retain the effect of receiving the true target class count during training.

**Table S6.** Deployment analysis. Each cell is mean ± sample SD followed by valid runs/4. Values are run-level equal-dataset summaries.

| Deployment metric | scOLAR | strict scBOL | Paired delta (scOLAR minus scBOL) |
|---|---|---|---|
| AUROC | 0.9726 ± 0.0012 [4/4] | 0.2741 ± 0.0591 [4/4] | 0.6985 ± 0.0597 [4/4] |
| AUPRC (AP) | 0.9871 ± 0.0006 [4/4] | 0.5660 ± 0.0262 [4/4] | 0.4210 ± 0.0266 [4/4] |
| Novel prevalence | 0.6564 ± 0.0018 [4/4] | 0.6564 ± 0.0018 [4/4] | — |
| Precision | 0.8837 ± 0.0052 [4/4] | 0.2500 ± 0.2516 [4/4] | 0.6338 ± 0.2483 [4/4] |
| Recall | 0.9782 ± 0.0015 [4/4] | 0.0430 ± 0.0319 [4/4] | 0.9352 ± 0.0323 [4/4] |
| F1 | 0.9284 ± 0.0023 [4/4] | 0.0664 ± 0.0477 [4/4] | 0.8621 ± 0.0469 [4/4] |
| Known retention | 0.7411 ± 0.0145 [4/4] | 1.0000 ± 0.0000 [4/4] | −0.2589 ± 0.0145 [4/4] |
| PredNovel fraction | 0.7273 ± 0.0062 [4/4] | 0.0288 ± 0.0209 [4/4] | 0.6985 ± 0.0233 [4/4] |
| All-PredNovel ARI | 0.3802 ± 0.0033 [4/4] | 0.0426 ± 0.0603 [2/4] | 0.5279 ± 0.3635 [2/4] |
| All-PredNovel AMI | 0.6607 ± 0.0028 [4/4] | 0.1885 ± 0.2667 [2/4] | 0.5822 ± 0.1654 [2/4] |
| All-PredNovel NMI | 0.6622 ± 0.0028 [4/4] | 0.1892 ± 0.2673 [2/4] | 0.5825 ± 0.1652 [2/4] |
| TP-only ARI | 0.3610 ± 0.0032 [4/4] | 0.0852 ± NA [1/4] | 0.7897 ± NA [1/4] |
| TP-only AMI | 0.6043 ± 0.0020 [4/4] | 0.3772 ± NA [1/4] | 0.4659 ± NA [1/4] |
| TP-only NMI | 0.6053 ± 0.0020 [4/4] | 0.3781 ± NA [1/4] | 0.4658 ± NA [1/4] |

*Note.* Undefined values remain missing and sample SD is NA when only one run is valid. Paired incomplete-case clustering deltas need not equal differences of marginal means. TP-only scores reuse the identical all-PredNovel Leiden partition and apply the ground-truth-novel mask only after clustering.

For strict scBOL, AUROC and average precision use `-source_max_logit` as a diagnostic ranking score. This derived score differs from the combined-head and remapping procedure that produces scBOL's native PredNovel decisions. Its ranking metrics are therefore supplementary diagnostics. Native precision, recall, F1, Known retention and PredNovel fraction describe the method's own operating point, with the true target class count supplied during training.

#### S5.3 End-to-end Known quantities

**Table S7.** Equal-dataset end-to-end Known summary across four independent runs. Values are mean ± sample SD. Every entry has $n = 4$.

| Method | Known retention | False-novel rate | Retained-known conditional accuracy | Joint correct-known rate |
|---|---|---|---|---|
| scOLAR | 0.7411 ± 0.0145 | 0.2589 ± 0.0145 | 0.9926 ± 0.0003 | 0.7359 ± 0.0141 |
| strict scBOL | 1.0000 ± 0.0000 | 0.0000 ± 0.0000 | 0.9584 ± 0.0064 | 0.9584 ± 0.0064 |

*Note.* Joint correct-known rate is a cellwise quantity and is not obtained by multiplying aggregate Known retention and retained-known conditional accuracy. Strict scBOL receives the true target class count during training.

Known classification under the common evaluator, deployment Known retention, accuracy conditional on retention and the joint correct-known rate measure different aspects of annotation. For scOLAR, most errors involving known cells arise when they are sent to the novel branch. Classification among the retained known cells remains accurate.

#### S5.4 Exploratory Quake Smart-seq2 metric analysis

We examined this dataset after observing the disagreement between aggregate recovery metrics. The analysis is descriptive and follows the primary evaluation. On Quake_Smart-seq2, scOLAR's Hungarian-matched Novel weighted accuracy is 0.6389 ± 0.0511, compared with 0.7172 ± 0.0374 for strict scBOL. Its Novel macro accuracy is 0.6854 ± 0.0649, compared with 0.5272 ± 0.0395. Local cluster purity, 15-nearest-neighbour agreement and same-label graph-edge agreement also remain high for scOLAR.

Numeric class 2 accounts for much of the weighted difference. It contains 4,394 cells, represents 30.33% of ground-truth-novel cells and contributes approximately −0.1114 to the four-run weighted-accuracy difference. Class 2 is a benchmark identifier with no inferred biological identity in this analysis. Its contribution explains the different directions of cell-weighted and class-balanced summaries, without establishing a causal mechanism.

#### S5.5 LCC scope

All 20 primary scOLAR run logs include LCC reports. LCC reuses each run's fixed partition of all PredNovel cells and produces a descriptive profile through CL ancestry. The Consistent designation records whether Top-3 coverage reaches 0.40. It does not indicate validated correctness or precision. The analysis covers a subset of communities and varies substantially across datasets, so it is not reported as an aggregate correctness endpoint.

### S6 Reproducibility

#### S6.1 Code and data availability

The scOLAR version 1.0.0 source code is available from GitHub at `https://github.com/Rijinna/scOLAR-project` and archived on Zenodo at `https://doi.org/10.5281/zenodo.22262387`. Both records identify the MIT license. Table S1 lists dataset sources and accessions. The standardised open-set partitions follow [5].

#### S6.2 Random seeds

The four primary runs use random seeds 101, 202, 303 and 404. Historical scBOL/scOLAR results use seeds 42, 123, 456 and 8888.

#### S6.3 Environment and computational cost

Table S8 gives the recorded software and hardware environment and development runtimes. Each recorded run uses one GPU. The retained records do not identify which GPU serves each dataset.

**Table S8.** Recorded software/hardware environment and retained development wall-clock times for the full model ($\lambda_{\text{HCL}} = 0.1$, 100 epochs). Intra-dataset entries preserve mean ± recorded spread across three development runs ($n = 3$). Replicate identities and the definition of the spread are unavailable in the retained records. Cross-dataset entries are single recorded values with no recorded replicate count.

| Environment | | Training time (min per run) | |
|---|---|---|---|
| Item | Value | Dataset | Time |
| Operating system | Debian GNU/Linux 12 (bookworm) | Cao | 3.1±0.4 |
| Python | 3.9.25 | Quake_10x | 5.2±0.2 |
| PyTorch / CUDA | 2.8.0 / 12.8 | Quake_Smart-seq2 | 3.9±0.4 |
| Scanpy / AnnData | 1.10.3 / 0.10.8 | Wagner | 2.9±0.1 |
| CL parsing | obonet 1.1.1, pronto 2.7.3 | Zeisel_2018 | 10.0±1.4 |
| GPU | NVIDIA RTX 3090, RTX 4090 | Mammary pair | 1.3 |
| CPU | Intel Xeon Platinum 8168, 96 cores | Pancreas pair | 2.6 |
| RAM | 251 GiB | Placenta pair | 21.8 |

*Note.* Times are end-to-end, including preprocessing, ontology construction, training, threshold calibration, inference and lineage-consistency analysis. The records include both GPU models but omit their dataset assignments. These development measurements provide timing context and cannot support a hardware-normalised runtime comparison.
